# Deep Learning-based Phenotype Imputation on Population-scale Biobank Data Increases Genetic Discoveries

**DOI:** 10.1101/2022.08.15.503991

**Authors:** Ulzee An, Ali Pazokitoroudi, Marcus Alvarez, Lianyun Huang, Silviu Bacanu, Andrew J. Schork, Kenneth Kendler, Päivi Pajukanta, Jonathan Flint, Noah Zaitlen, Na Cai, Andy Dahl, Sriram Sankararaman

**Affiliations:** ComputerScience Department, UCLA, Los Angeles, CA, USA; Department of Human Genetics, David Geffen School of Medicine at UCLA, Los Angeles, CA, USA; Helmholtz Pioneer Campus, Helmholtz Zentrum München, Neuherberg, Germany; Computational Health Centre, Helmholtz Zentrum München, Neuherberg, Germany; School of Medicine, Technical University of Munich, Munich, Germany; Virginia Institute for Psychiatric and Behavioral Genetics and Department of Psychiatry, Virginia Commonwealth University, Richmond, VA, USA; Institute of Biological Psychiatry, Mental Health Center - Sct Hans, Copenhagen University Hospital, Copenhagen, Denmark; Neurogenomics Division, The Translational Genomics Research Institute (TGEN), Phoenix, AZ, USA; Section for Geogenetics, GLOBE Institute, Faculty of Health and Medical Sciences, Copenhagen University; Institute for Precision Health, David Geffen School of Medicine at UCLA, Los Angeles, CA, USA; Neurology Department, UCLA, Los Angeles, CA, USA; Section of Genetic Medicine, University of Chicago, Chicago, IL, USA; Department of Computational Medicine, UCLA, Los Angeles, CA, USA

## Abstract

Biobanks that collect deep phenotypic and genomic data across large numbers of individuals have emerged as a key resource for human genetic research. However, phenotypes acquired as part of Biobanks are often missing across many individuals, limiting the utility of these datasets. The ability to accurately impute or “fill-in” missing phenotypes is critical to harness the power of population-scale Biobank datasets. We propose AutoComplete, a deep learning-based imputation method which can accurately impute missing phenotypes in population-scale Biobank datasets. When applied to collections of phenotypes measured across ≈ 300K individuals from the UK Biobank, AutoComplete improved imputation accuracy over existing 2 methods (average improvement in *r*^2^ of 18% for all phenotypes and 42% for binary phenotypes). We explored the utility of phenotype imputation for improving the power of genome-wide association studies (GWAS) by applying our method to a group of five clinically relevant traits with an average missigness rate of 83% (67% to 94%) leading to an an increase in effective sample size of ≈2-fold on average (0.5 to 3.3-fold across the phenotypes). GWAS on the resulting imputed phenotypes led to an increase in the total number of loci significantly associated to the traits from four to 129. Our results demonstrate the utility of deep-learning based imputation to increase power for genetic discoveries in existing biobank data sets.

## Introduction

The past decade has seen the growth of datasets that collect deep phenotypic and genomic data across large numbers of individuals. While these population-scale biobanks aim to capture a wide range of phenotypes across the population (including demographic information, laboratory tests, imaging, medication usage, and diagnostic codes), phenotypes in this setting are frequently missing across many of the individuals for reasons such as cost or difficulty of acquisition *(e.g.,* phenotypes derived from imaging scans and other potentially invasive procedures). As a result, our ability to study clinically-relevant phenotypes or diseases using biobank data remains limited.

The ubiquity of missing data in the biomedical domain has motivated extensive work into statistical methods for imputing or “filling-in” missing data [1,2,7,8,9,37,50] (see Section S1 for additional related work). Accurate imputation of large numbers of phenotypes and individuals in population-scale Biobank data presents several challenges. First, accurate imputation requires faithfully modeling the dependencies across the phenotypes. Such dependencies can arise due to genetic or environmental effects that are shared across phenotypes. Accumulating evidence for the abundance of shared genetic effects (pleiotropy) even amongst seemingly unrelated phenotypes suggests that the ability to model dependencies across large numbers of collected phenotypes could substantially improve imputation accuracy. Second, patterns of missingness in these datasets tend to be complex (for example, individuals that were not administered a questionnaire will be missing for all answers relevant to the questionnaire) so that an imputation method would need to be able to use the relevant observed entries to impute the missing entries. Third, the method needs to be scalable. Thus, imputation methods that can accurately impute phenotype in the presence of complex patterns of missingness while being scalable are needed.

Here we propose AutoComplete, a deep-learning method based on an auto-encoder architecture designed for highly incomplete biobank-scale phenotype data. Our use of deep learning for imputation is motivated by the ability of neural networks to learn potentially complex dependencies among phenotypes (as shown in the application of neural networks to other biological datasets [60, 61, 62, 63, 64]). Earlier works, however, have relied on access to individuals with no missing phenotypes to learn the imputation model [43] (such an approach would substantially reduce the data available to learn model) or have assumed that entries in a dataset are missing completely at random[34, 41]. To be able to impute in the presence of realistic patterns of missingness, we employed copy-masking, a procedure that propagates missingness patterns present in the data [37]. AutoComplete can impute both binary and continuous phenotypes while scaling with ease to datasets with half a million individuals and millions of entries.

We compared the accuracy of AutoComplete to state-of-the-art missing data imputation methods on two collections of phenotypes derived from the UK Biobank (UKBB) [5]: a set of 230 cardiometabolic-related phenotypes and a set of 372 phenotypes from an on-going study of psychiatric disorders, each measured across ≈ 300, 000 unrelated white British individuals. AutoComplete improved squared Pearson correlation (*r*^2^) by 18% on average over the next best method (SoftImpute [9]) and 42% on average for binary phenotypes. We then explored the utility of our method in increasing the power to detect genetic associations for traits of interest. We demonstrated the validity of this procedure by simulating missingness in fully observed phenotypes and showing that all significantly associated loci identified by GWAS on imputed phenotypes were associated with the originally observed phenotypes. We then performed genome-wide association studies (GWAS) of five clinically relevant but highly missing phenotypes (missingness rate of 83% on average and range 67%~94%) from the UKBB with the aid of imputed values using AutoComplete. We observed an increase in effective sample size of 1.96-fold on average (0.5-fold to 3.3-fold across the phenotypes). GWAS on the resulting imputed phenotypes yielded a total of 129 significantly associated loci (compared to four loci significantly associated with the original phenotypes) with a mean increase of 25 loci across the tested phenotypes. AutoComplete can scale to ≈ 300, 000 individuals and ≈ 400 phenotypes from the UKBB with empirical run times of ~ 6 hours. Our results illustrate the value of deep-learning based imputation for genomic discovery.

## Results

### Methods overview

AutoComplete is based on an autoencoder (a type of neural-network) that is capable of simultaneously imputing continuous and binary-valued features. Given a vector of features that represent the phenotypes measured on an individual (some of which might be missing), AutoComplete maps the features to a hidden-representation using a non-linear transformation (encoder) which is then mapped back to the original space of features to reconstruct the phenotypes (decoder). In this process, AutoComplete imputes missing phenotypes (Figure 1).

**Figure 1:**
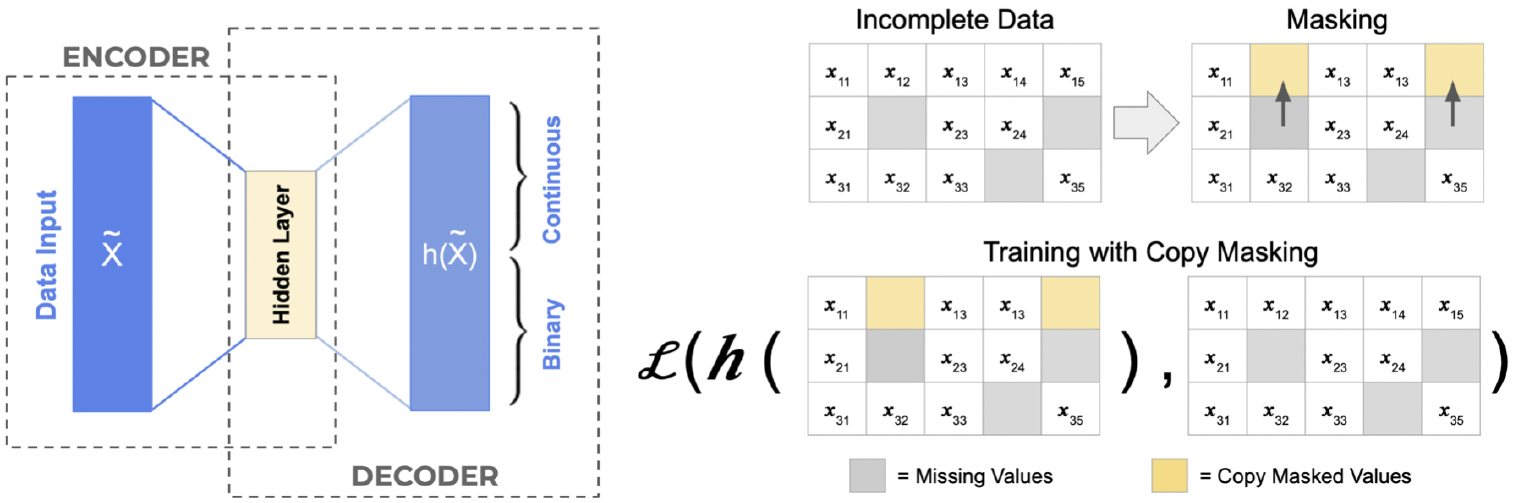
AutoComplete architecture. AutoComplete defines a feed-forward encoder-decoder architecture *h* trained using copy masking -- a procedure that simulates realistic missingness patterns that the model uses to impute missing values. AutoComplete minimizes the loss function *L* that is defined over the observed and masked values. AutoComplete supports the imputation of continuous and binary features.

AutoComplete aims to learn the autoencoder by masking features that are originally observed in the data and searching for the parameters of the autoencoder that can reconstruct the masked and observed features with minimal error. To enable AutoComplete to impute in the presence of realistic missingness patterns, we employed copy-masking, a procedure which propagates missingness patterns already present in the data [37].

### Experiment overview

We evaluated the accuracy of phenotypes imputed by AutoComplete on two collections of UKBB phenotypes: a set of 230 cardiometabolic phenotypes derived from patient records and imaging data, and a larger set of 372 phenotypes from an on-going study on Major Depressive Disorder (MDD). Each collection contains phenotypes measured across ≈ 300, 000 unrelated individuals of white British ancestry, where phenotypic entries were, on average, 49% and 46% missing respectively (Table S1).

We compared the accuracy of AutoComplete to a representative selection of imputation methods that could be applied at scale. We considered K-Nearest Neighbors (KNN), missForest [6], and MICE [7], among the most widely-used imputation methods routinely available in data science packages [8]. We also considered SoftImpute [9] based on its consistently high imputation accuracy in prior works [11,37]. Finally, we also evaluated two generative deep-learning methods: a generative-adversarial imputation method, GAIN, [10] and a deep generative model, HI-VAE, [11] (see Section S1 for a more detailed description of related methods).

In determining which methods scale, we assessed empirical runtimes to fit each imputation method for increasingly large subsets of the psychiatric disorders dataset to determine that missForest and MICE do not efficiently scale to the UKBB datasets leading us to exclude missForest and MICE from our large-scale comparisons (Figure S1).

To quantify the accuracy of each method on previously unseen individuals, we adopt a train-test split (50% each) of the two datasets such that all hyperparameter tuning and training were performed on the training set while evaluations of all methods were performed on the test set (see Methods Section).

To evaluate the imputation methods, we simulated missing entries by masking originally 2 observed phenotypes across a range of missingness levels (1% ~ 50%). We examined *r*^2^ between imputed and originally observed values as the primary metric, given its compatibility with continuous and binary phenotypes and its interpretation in terms of the effective sample size [12]. We examined imputation accuracy on binary phenotypes separately using both *r*^2^, area under the precision-recall curve (AUPR) and the area under receiver operating characteristic curve (AUROC). For each metric, we quantified standard error and confidence intervals using bootstrapping of 200 replicates. To test for significant differences in the imputation accuracy obtained by each method, we performed a two-tailed significance test of the difference in means using the bootstrap standard errors.

In addition to the two larger collections of phenotypes, we evaluated our method for a smaller subset of the cardiometabolic dataset consisting of 86 phenotypes, allowing comparisons with MissForest [6] and MICE Forest [2] in tractable running times (Section S2).

### AutoComplete significantly improves imputation accuracy

AutoComplete obtained the most accurate imputations across all levels of missingness (from 1% to 50%) in the tested datasets (Table 1, Figure 2). Imputation accuracy was generally higher in the cardiometabolic dataset for all methods relative to the psychiatric disorders dataset which we hypothesize can be attributed, in part, to the greater proportion of missing entries in the latter (Table S1). Further, the imputation accuracy of all methods decreased with increasing levels of missingness. While SoftImpute (based on a linear model) was the most accurate among existing methods, AutoComplete obtained the the highest overall accuracy (an improvement of 7% and 25% for cardiometabolic and psychiatric disorder phenotypes over SoftImpute, respectively) indicating the value of modeling non-linear relationships among phenotypes (Figure 2).

**Table 1:**
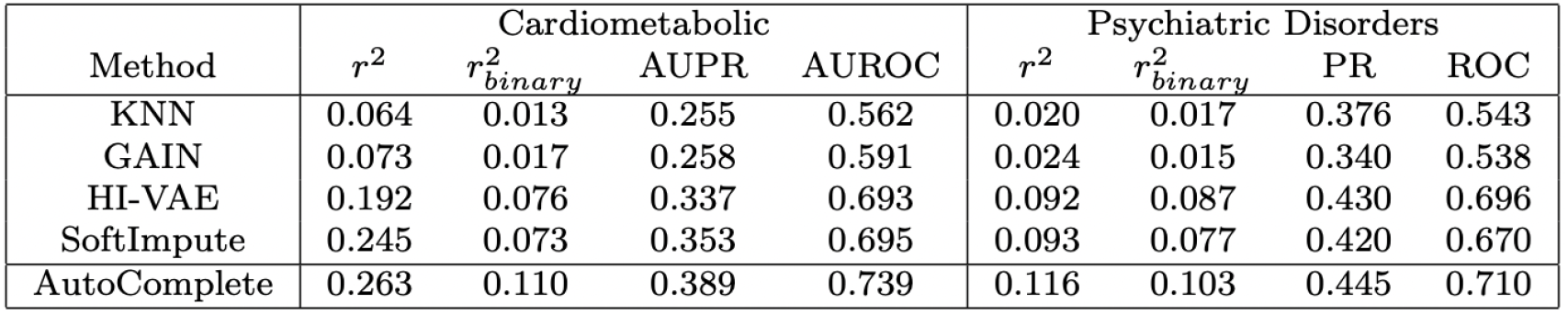
Summary of imputation accuracy. Average accuracy in Pearson’s *r^2^* (for all phenotypes and for binary phenotypes), area under the precision-recall curve (AUPR), and area under the receiver operating characteristic curve (AUROC) across all simulations of 1%~50% missing data.

**Figure 2:**
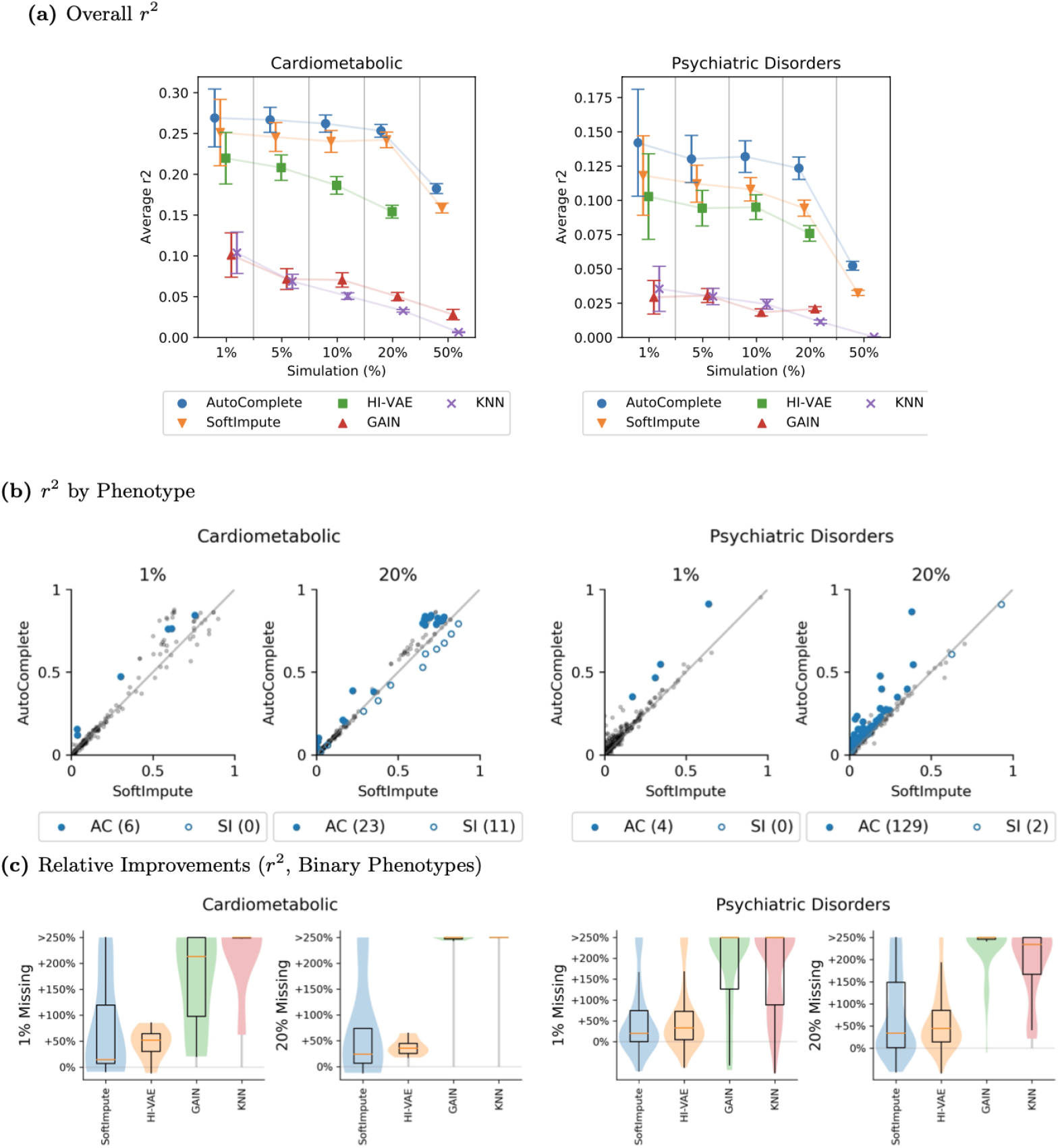
AutoComplete provides accurate imputations across a range of simulation settings. (a) Overall imputation accuracy across a range (1%~50%) of simulated missingness. We report the average *r^2^* across phenotypes in each dataset (bars denote average of 95% CIs). Only AutoComplete and SoftImpute could be reliably fit for all levels of simulated missingness (HI-VAE and GAIN could not be evaluated for higher levels of simulated missingness due to instabilities in training). (b) Imputation accuracy (Pearson’s *r^2^*) for individual phenotypes were compared between AutoComplete and SoftImpute (next-best). Accuracy was measured under simulated levels of missing data, highlighted for settings of 1%~20%. Bold dots indicate a significant difference in accuracy (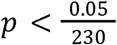 and 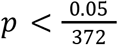 respectively for cardiometabolic and psychiatric disorder phenotypes). (c) Relative improvements in *r^2^* imputation accuracy for binary-valued phenotypes. Comparisons were aggregated over phenotypes between AutoComplete and each existing method.

AutoComplete significantly improved *r*^2^ for 6 (23) phenotypes over SoftImpute with 1% (20%) missingness in the cardiometabolic dataset (Table S2; *p*-value 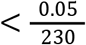 correcting for the 230 phenotypes tested). Analogously, AutoComplete significantly improved *r*^2^ for 4 (129) phenotypes with 1% (20%) missingness in the psychiatric disorders dataset (Table S2; *p*-value 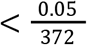 correcting for the number of phenotypes tested). Among the most significantly improved phenotypes were report of statin use (56%) and biomarker measurements from full-body DXA scans [13] such as arm lean mass (38%), recurrent MDD (59%) [51] and receiving medication for psychotic condition (107%).

The improvements in imputation accuracy were particularly substantial for binary phenotypes. Here, AutoComplete obtained a relative improvement over the next best method (SoftImpute) of 56% in *r*^2^ on the cardiometabolic data and 34% on the psychiatric disorders data across all simulations (Figure 2(c)). We found qualitatively similar trends for other metrics such as the area under the precision-recall curve (AUPR) and area under the ROC curve (AUROC) for the imputations obtained on the binary phenotypes (Figure S2; Table S2). In comparison to SoftImpute, AutoComplete imputation obtained a relative increase in AUPR of 11% and AUROC of 7% in the cardiometabolic dataset and an increase of 6% for both metrics in the psychiatric disorders dataset (Table 1).

We performed a separate experiment on a small-scale subset of UKBB where we compare AutoComplete to methods that include missForest and MICE that could not scale to the full UKBB phenotypes and find that AutoComplete remains the most accurate method in this setting (Section S2, Figure S5).

Finally, we also explored the importance of the copy-masking procedure. We compared AutoComplete trained with copy-masking to training the denoising autoencoder with masking performed uniformly at random (Section S3). For the setting of 1% missing values, the highest average *r*^2^ obtained through uniformly random masking was 0.121 in comparison to 0.142 with AutoComplete (17%) with similar trends in tests with increasing missingness (average 16% improvement using copy-masking; Section S3, Figure S6). We further assessed the importance of copy-masking in the evaluation step to simulate missing values to measure imputation accuracy. Instead of copying existing missing patterns, we chose values to be missing uniformly randomly among all observed values until 1%~50% of the observed data was withheld for imputation in the psychiatric disorders dataset (SectionS3, Figure S7). When not propagating the structured missingness for testing, the imputation accuracy (*r*^2^) of AutoComplete was inflated to 0.213 on average (0.116 originally) while imputation accuracy of LifetimeMDD grew to 0.987 (0.468 originally). Softimpute also behaved similarly in this setting. We therefore conclude that copy-masking is integral to evaluating imputation accuracy and that AutoComplete benefits from mimicking realistic missingness patterns that aid the denoising behavior of the deep learning model.

### Imputed phenotypes improve the power to detect novel genetic risk factors

We explored the utility of phenotypes imputed using AutoComplete for improving power in genome-wide association studies (GWAS). We first examined the reliability of a GWAS performed on imputed phenotypes through a simulation in which we introduced missingness into a phenotype that has a low level of missingness in the psychiatric disorder dataset (alcohol consumption which is < 0.1% missing). The phenotype was masked in 67% of the samples such that the sample size available for a GWAS was *N* = 114, 778. AutoComplete was then applied to impute the masked entries, recovering the original sample size of *N* = 337,126 with an imputation accuracy (*r*^2^) of 0. 397.

We performed GWAS, in turn, on the observed phenotypes restricted to 33% of the individuals (*N* = 114, 778) and on the imputed phenotypes on the full sample size of *N* = 337,126 (see Methods). Using the AutoComplete-impute phenotypes increases the number of significantly associated SNPs (p < 5 × 10) from 14 to 87 with a corresponding increase in the number of associated loci from three to six (Figure 3(a), QQ-plot of GWAS with imputed phenotype in Figure 3(b)).

**Figure 3:**
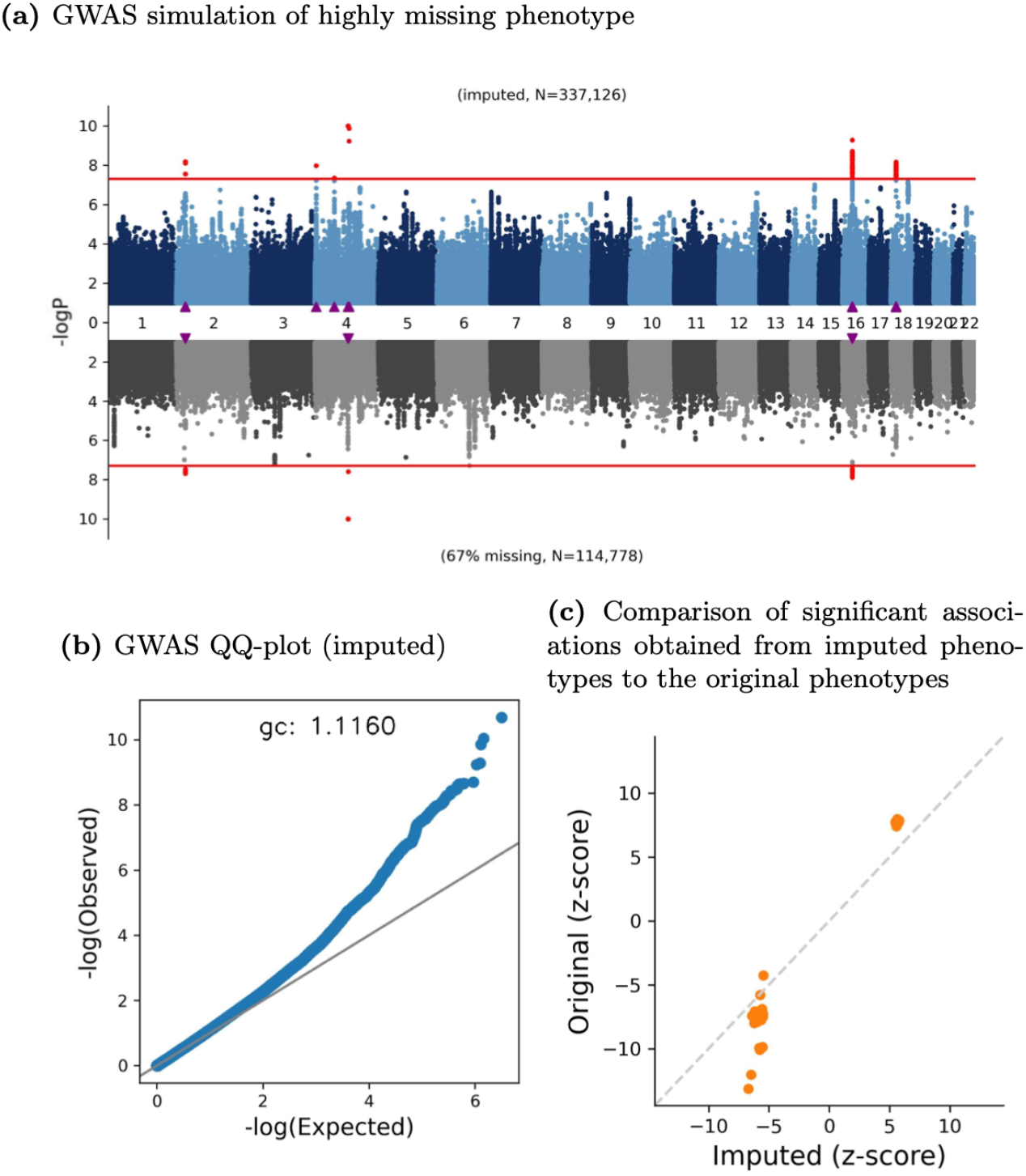
AutoComplete imputation leads to replicable associations. We simulated missing values (67% missingness) in a phenotype (Alcohol consumption) that had low missingness and imputed missing values using AutoComplete. (a) GWAS for imputed alcohol consumption (top, N=337,126) compared to GWAS on the observed phenotype (bottom, N=112,297). Triangle markers indicate significantly associated loci (significantly associated SNPs in red, *p* < 5 × 10 threshold indicated by red horizontal lines). The number of significant loci increased from three to six corresponding to an increase in the number of significantly associated SNPs from 14 to 87. (b) The corresponding QQ-plot of the GWAS augmented with imputed measurements is shown. (c) We attempted to test whether the significant associations obtained from the imputed phenotypes (“Imputed”) were truly associated with the original phenotype (100% observed in data, “Original”). Z-scores of effect sizes for each experiment were plotted _8 against one another. Significant associations in the imputed phenotype (*p* < 5 × 10^−8^) were determined as having a verifiable effect in the fully observed phenotype GWAS for *p* < 5 × 10^−4^ (accounting for the number of SNPs tested). All 87 SNPs that were significantly associated in the imputed test could be verified as being strongly associated in the original test.

We tested the association of each of the 87 significantly associated SNPs with the observed phenotype in the full sample and found that all 87 significantly associated SNPs were found to be significant (*p* < 5 × 10 accounting for the number of hypotheses tested; Figure 3 (c)). Further, we compared the marginal effects of SNPs estimated from GWAS on each of the sub-sampled phenotypes and the imputed phenotype to the phenotype observed in the full sample and observed that the squared Pearson correlation between the effect sizes of SNP associations increases from 0. 35 (in line with expectation that the subsampled GWAS was obtained on a 33% subset of individuals from the full sample [70]) in the subsampled GWAS to 0. 6 in the imputed GWAS.

We repeated the simulation and evaluation procedure for an additional phenotype with low-levels of missingness (insomnia; Figure S3(a), QQ plot in Figure S3(b)) and found qualitatively similar results. The imputed phenotypes provided an effective increase in power (imputation accuracy of *r*^2^ =0.183) with an increase from 10 to 28 significantly associated SNPs corresponding to an increase in 3 to 6 significant loci. We found that 26 of the 28 associated SNPs were found to be associated in the original set of observed phenotypes *(p* < 5 × 10 accounting for the number of hypotheses tested; Figure S3(c)).

We then performed genome-wide association studies (GWAS) for five phenotypes (LifetimeMDD [51], anxiety with drug use, using medication for MDD, number of MDD episodes, and having ever taken cannabis) which were clinically relevant to Major Depressive Disorder (MDD) in the psychiatric disorder dataset after using AutoComplete to impute all missing values. To verify that these phenotypes are accurately imputed, we confirmed that the ratio of variance of the imputed observations to the original observations (analogous to the metrics used to measure the quality of genotype imputation [65, 66]) is sufficiently large (> 0.2 across these phenotypes; we had 62 phenotypes pass this threshold in the psychiatric disorders dataset).

We estimated the effective gain in sample size resulting from this procedure for each phenotype (Table S3; see Methods). We observed an increase in the effective sample size of 1.96-fold on average: LifetimeMDD had an effective sample size of 193,379 from 67,164 originally observed samples (a 1.88-fold increase) while the increase was lowest for Cannabis Ever Taken (110,188 to 164,653 resulting in a 0.5-fold increase).

We performed GWAS with the imputed phenotypes and observed an increase in the total number of significantly associated loci across all phenotypes of interest from four (when analyzing the original phenotypes) to 129, with the mean increase being 25 loci (range 13 to 37; Table 2). The GWAS results of cannabis ever taken (from 2 loci to 28 loci) and anxiety with drug use (from 0 to 16 loci) are shown in Figures 4(a) and 4(b) respectively (see QQ-plots in Figure S4).

**Table 2:** Comparison of GWAS performed on five phenotypes of interest with high levels of missingness before and after imputation. We report the number of significantly associated loci when performing GWAS on the original phenotype in comparison to the number of significant loci obtained when performing GWAS after imputing the missing phenotypes using AutoComplete. The number of additionally discovered loci (More Loci) in applying AutoComplete were tallied in comparison to original phenotype without imputation.

| Phenotype | Missing | Original<br># Loci | AutoComplete<br># Loci | More Loci |
| --- | --- | --- | --- | --- |
| LifetimeMDD | 80% | 1 | 34 | 33 |
| Anxiety with Drug Use | 95% | 0 | 16 | 16 |
| Medication for MDD | 91% | 1 | 14 | 13 |
| MDD Episodes | 83% | 0 | 37 | 37 |
| Cannabis Ever Taken | 67% | 2 | 28 | 26 |

**Figure 4:**
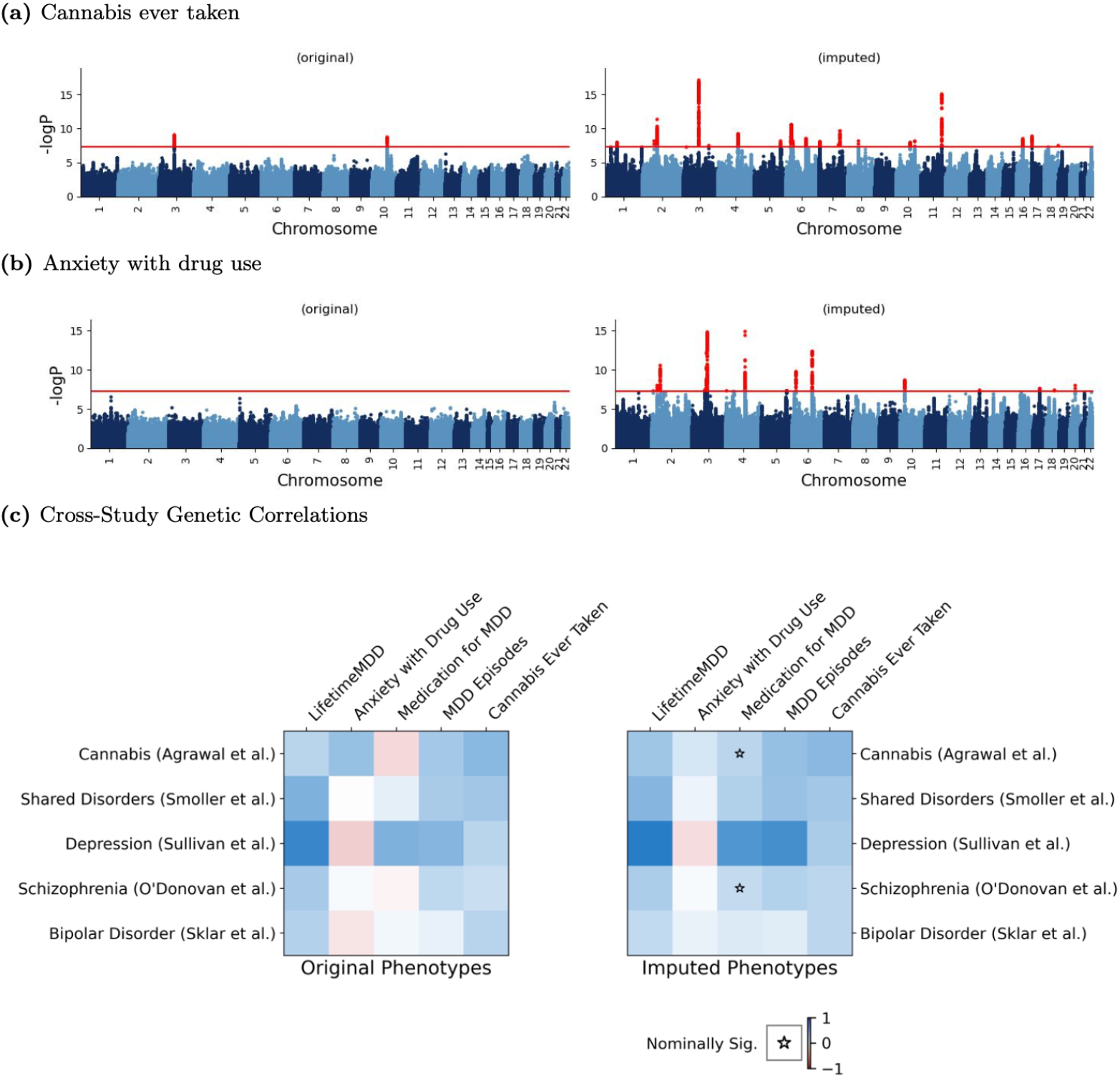
GWAS on AutoComplete imputation phenotypes lead to additional significant genetic associations. For a select group of highly missing phenotypes which are clinically relevant to psychiatric disorders, the results of Genome-Wide Association Study (GWAS) were compared between limiting the analysis to only the observed phenotypes and including measurements imputed using AutoComplete for all individuals. An increase in the number of significant associations were observed for (a) cannabis ever taken, (b) anxiety with drug use, and several more phenotypes (significant associations in 8 red, *p* < 5 × 10^−8^ threshold indicated by the red horizontal line). (c) The genetic correlation (*r_G_*) was assessed between the GWAS of highly missing phenotypes originating from the UK Biobank and out-of-sample GWAS of similar traits (left). Colors indicate *r_G_* closer to 1 (blue) or −1 (red). The genetic correlation was computed again after imputing all missing phenotypes with AutoComplete (right). We tested for a significant change in genetic correlations from using imputed phenotypes and non-imputed phenotypes. Stars indicate nominally significant change (*p* < 0. 05). None of the genetic correlations significantly changed between the two settings after correcting for multiple hypothesis testing (Bonferroni correction, 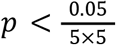.

We investigated the relationship between the original and the imputed phenotypes by estimating the genetic correlation (*r_G_*) between each of these phenotypes (both the original and the AutoComplete-imputed phenotypes) and phenotypes related to psychiatric disorders obtained from cohorts outside the UK Biobank cohort: cannabis-use disorder [14], multiple psychiatric disorders [15], MDD [16], schizophrenia [17], and bipolar disorder [18]. Genetic correlation was estimated using LDSC for all pairs of 5 external phenotypes and 5 UK Biobank MDD phenotypes [19,20]. We compared the results from observed UK Biobank phenotypes before imputation and those from Autocomplete imputed phenotypes (Figure 4(c)). We observed that the genetic correlations remained largely similar between using the observed and imputed UK Biobank phenotypes (p-value < 0.05/25 accounting for 25 comparisons between the 5 phenotypes and 5 external studies). Thus, the genetic component of the phenotypes imputed by AutoComplete tend to be qualitatively similar to the original observed phenotypes.

## Discussion

The ubiquity of missing data in population-scale biobanks necessitate effective methods for imputation. Here, we describe AutoComplete, a deep-learning approach to imputation which we demonstrate to be accurate and efficient for imputing phenotypes in the UK Biobank.

AutoComplete increased imputation accuracy of highly missing phenotypes related to cardiometabolic and psychiatric disorders in comparison to state-of-the-art linear methods. This implies that understanding nonlinear dependencies among phenotypes in biobank data is important. Patterns of missingness are often structured for biobank-type data as a consequence of the data-gathering procedures. We also observed that realistic simulations of missing data make a substantial contribution to the accuracy of the model learned for imputation (Section S3). Our use of copy-masking provides a straightforward and general approach for training deep-learning methods in the presence of complex, structured missingness thereby broadening their applicability.

We discuss limitations of our method and directions for future work. First, the basic autoencoder architecture underlying our method can be extended in many ways. While we determined through cross validation that the majority of the imputation accuracy is gained architecturally from the first three layers and the support for continuous and binary imputations, a fuller exploration of the architecture of the neural network could lead to further improvements to accuracy. Second, as biobanks collect diverse data modalities, including imaging, time-series and multi-omic data, imputing missing data that arises in the context of these diverse data types remains a challenge. The modularity of the underlying neural network architecture will enable our method to deal with the diversity of phenotypic data types that are being gathered and we leave this as a promising direction for future work. Third, our method does not provide estimates of uncertainty in the imputed values. Propagating imputation uncertainty in downstream analysis could, in principle, lead to more robust inferences. The typical approach to estimate uncertainty uses multiple imputation in which multiple imputed datasets are constructed by sampling from the predictive distribution (assuming a Bayesian model). However, multiple imputation strategies tend to be challenging to apply to large-scale datasets. While our current results suggest that our imputed phenotypes yield robust genetic associations, reporting of uncertainty in the imputed phenotypes is an important direction for future work. Finally, the consequence of using a deep-learning method is that the resulting imputation phenotypes are often challenging to interpret. Such interpretations are critical to understanding whether an imputed phenotype is enriched for the genetic component of the original phenotype. Methods for interpreting deep learning methods is an area of active research [54, 23] and could be extended to our setting. Analyzing the signals driving our imputation method when applied to biological datasets could reveal distinct subtypes of a disease and could provide insights into disease etiology. Interpretable components could also give higher credence to the imputed phenotypes.

## Methods

### Datasets

The UK Biobank Dataset (UKBB) [5] makes available genetic data for up to half-million individuals and thousands of traits. We gathered two collections of phenotypes in the UKBB which were analyzed together from existing studies.

We collected a group of 230 cardiometabolic phenotypes [13,24]. These consisted of phenotypes and serum biomarkers derived from body imaging and laboratory measurements relevant to cardiometabolic disorders, consumption of prescribed drugs (e.g. medication for cholesterol or aspirin), measures of daily physical activity and food consumption, as well as anthropometric and general demographic information. In addition, we collected ICD10 and ICD9 codes relating to non-alcoholic fatty liver disease (NAFLD) [55, 56] and ICD10, ICD9, and OPCS-4 codes relating to coronary artery disease (CAD) following [67].

We constructed a second dataset of 372 phenotypes related to psychiatric disorders. This included lifetime and current MDD symptom screens [68,69], psychosocial factors, comorbidities, family history of common diseases, a broad range of demographic information, as well as both deep and shallow definitions of MDD derived from symptom questionnaires using clinical diagnostic criteria or self-reports [51]. Both datasets consist of ≈ 300, 000 white British unrelated individuals. Each of these collections included a mix of continuous and binary-valued phenotypes (Table S1). Missingness rates for phenotypes across individuals varied from 0% (age, sex) and up to 99% (addiction, self-harm).

For each dataset containing *N* individuals and *P* phenotypes, a data matrix of dimension *N* × *P* was created including missing values. Approximately ≈ 50% of all individuals were reserved for testing (evaluating the accuracy of the methods) while the remainder was used for training and any hyperparameter tuning for all methods. Continuous phenotypes were normalized to have zero mean with unit variance per phenotype. Binary-valued phenotypes were processed specific to the capabilities of each method; for methods which did not handle binary data, labels were converted from 0,1 to – 0. 5, 0. 5 and treated as continuous values. To prevent information leakage, statistics of the training split were used to normalize the test split.

### AutoComplete

AutoComplete is based on a type of neural-network that is capable of simultaneously imputing continuous and binary-valued phenotypes. For each individual, AutoComplete considers a fixed list of phenotypes including missing values and reconstructs all phenotypes from a latent representation using an auto-encoder architecture. Of the input phenotypes, missing entries were masked (set to zero), then all observed phenotype values were transformed to a hidden representation in the encoding stage. The decoding stage transforms the hidden representation back to the input space such that all phenotypes were reconstructed. To support heterogeneous data types, imputed entries corresponding to binary phenotypes were obtained as the output of a sigmoid function so that these entries lie in the range [0,1].

Let 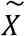 denote a *N × P* phenotype matrix such that 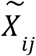 is the value of j-th phenotype measured on i-th individual, *M* denote a *N × P* indicator matrix (termed the Mask matrix) where *M_ij_* =1 if the j-th phenotype is observed for i-th individual and *M_ij_* = 0 otherwise. For simplicity, continuous and binary phenotypes were organized in 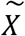 such that the first *C* phenotypes were continuous.

*h* denotes the nonlinear function corresponding to the autoencoder. The function *h* imputes both missing phenotype values and reconstructs observed ones. During imputation, only the imputed missing values are used. Using the LeakyReLU function Φ as a nonlinearity in the hidden layer and the sigmoid function s which was applied to binary-valued imputations, we define for the case of 1-hidden layer the following feed-forward function *h* (additional hidden layers could be defined analogously):

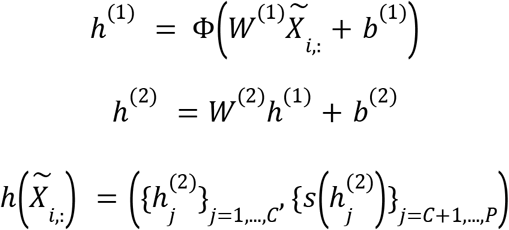

where

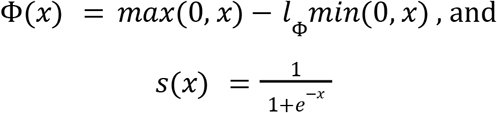

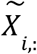 denotes row *i* of 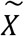 (equivalently the vector of phenotypes associated with individual *i*)

For each layer, the learnable weight parameter *W* is a *D* × *P* matrix where D is the dimension of the hidden representation while the bias vector *b* is of length *D*.

Given function h, the final imputed matrix 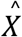 is constructed from 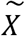 as follows:

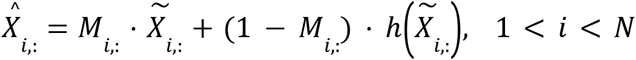

Here. denotes entrywise product.

In training, we promoted imputation using *h* such that both truly observed and masked phenotype values were subject equally to a reconstruction loss. Observed values were withheld based on existing missingness patterns which were randomly drawn from the dataset by individuals, then applied to other individuals; a process we refer to as copy-masking. To do this, a binary mask vector 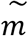 is drawn from the rows of the mask matrix *M* and was applied to the input of *h* such that for individual-*i*, the *j*-th phenotype would be masked when 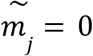 or unmodified when 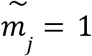. We controlled the prevalence of masking in training by the parameter *ρ* which was the probability one individual would receive a copy-mask. The masking process of AutoComplete is illustrated in Figure 1.

A joint loss function was defined over observed and masked values such that Mean Square Error and Cross Entropy loss were applied to continuous and binary phenotypes respectively. For simplicity the two types of phenotypes were partitioned by index C. The joint loss function was applied over all values which were originally observed:

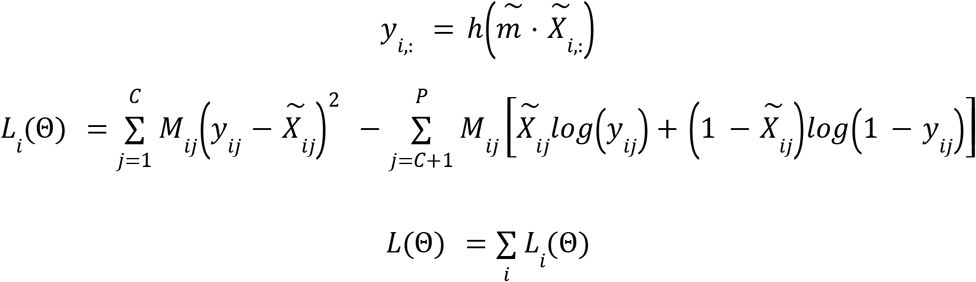

The parameters 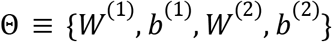 of *h* were optimized with respect to the objective *L*. Stochastic Gradient Descent (SGD) [25] was used to fit the neural net, where the initial learning rate, momentum, and mini-batch size were also determined using cross-validation. The weights and biases of the network were initialized using the Kaiming Uniform distribution, and the slope parameter of LeakyReLU was initialized as *i*_Φ_ = 0. 01. Training proceeded up to 500 epochs or convergence based on the validation split of the training split. A single RTX8000 GPU was used for all experiments.

#### Copy Masking

We implemented copy-masking, a simulation procedure to induce realistic patterns of missingness on observed data. For a given individual, missingness patterns from other random individuals were applied with probability *ρ*. This approach strives to maintain the realistic missingness patterns in datasets while introducing simulated missing values. Observed values were withheld based on existing missingness patterns which were randomly drawn from the dataset by individual, then applied to other individuals. Therefore, a binary vector referred to as 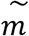 of length P where 1s indicated non-missing entries was drawn randomly uniformly from the rows of the mask matrix *M*, then applied to the input of h such that for individual-*i*, the *j*-th phenotype could be masked as 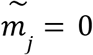 or unmodified as 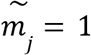. We controlled the prevalence of masking in training by the parameter *ρ* which was the probability one individual would receive a copy-mask.

In contrast, uniform randomly withholding observed values could distort the distribution of the features, *e.g.,* when two features have correlated missingness. In training, we used copy-masking to simulate missing values such that they were reconstructed by the autoencoder. For testing, copy-masking was used to to introduce additional missingness to the original data in the range of 1% ~ 50%, withholding the masked observed values for evaluation.

### Hyperparameter Tuning

Hyper-parameter search was performed for SoftImpute using cross-validation on the training set, adjusting the nuclear norm and maximum rank until reconstruction mean-squared error was minimized. HI-VAE requires choosing *y* hidden units per phenotype, the size of the latent dimension, *z*, and the number of latent mixture components *s*. The settings of *y* = 10, *z* = 8, *s* = 1 were chosen as additional mixtures did not appear to improve imputation in cross-validation. GAIN was configured with a hint rate of 0.9 and the weighting of the mask reconstruction term was chosen as the default *α* = 10. The generator and discriminator architectures of GAIN each consisted of 3 fully-connected layers and ReLU as non-linearities. Similarly to MissForest and MICE, KNN had difficulty scaling to the biobank data, and was evaluated once with *K* = 10.

Several hyperparameters were tuned using cross validation for AutoComplete in optimizing the objective function. The chosen hyperparameters were learning rate of 2. 0 and momentum of 0.9 for SGD optimization, learning rate decay of 0.5 for 5 epochs of non-improvement, batch size of 4096, and leaky ReLU parameter of 0.01. For the cardiometabolic and psychiatric disorders datasets, the encoding dimension *D = P* (no reduction), hidden layer count of 1, and masking amount ρ = 30% were determined through cross-validation using the training set.

### Details of GWAS analysis

We used imputed genotypes available from the UKBB for the individuals that were included in the phenotype imputation. We performed stringent filtering on the imputed variants, removing all insertions and deletions (INDELs) and multi-allelic SNPs: we hard-called genotypes from imputed dosages at 9,720,420 biallelic SNPs with imputation INFO score greater than 0.9, MAF greater than 0.1%, and p-value for violation of Hardy-Weinberg equilibrium > 10^-6^, in individuals with a genotype probability threshold of 0.9 (individuals with genotype probabilities below 0.9 would be assigned a missing genotype). Of these, 5,776,313 SNPs are common (MAF > 5%). We consistently use these SNPs for all analyses in this study.

We used 20 PCs computed with FlashPCA [52] on 337,126 White-British individuals in UKBB and genotyping arrays as covariates for all GWAS. We performed principal component analysis (PCA) on directly genotyped SNPs from samples in UKBB and used PCs as covariates in all our analyses to control for population structure. From the array genotype data, we first removed all samples who did not pass QC, leaving 337,126 White-British, unrelated samples. We then removed SNPs not included in the phasing and imputation and retained those with minor allele frequencies (MAF) ≥ 0.1%, and P value for violation of Hardy-Weinberg equilibrium > 10^-6^, leaving 593,300 SNPs. We then removed 20,567 SNPs that are in known structural variants (SVs) and the major histocompatibility complex (MHC) as recommended by UKBB [5], leaving 572,733 SNPs. Of these, 334,702 are common (MAF > 5%), and from these common SNPs we further filtered based on missingness < 0. 02 and pairwise LD *r*^2^ < 0.1 with SNPs in a sliding window of 1, 000 SNPs to obtain 68,619 LD-pruned SNPs for computing PCs using FlashPCA. We obtained 20 PCs, their eigenvalues, loadings and variance explained, and consistently use these PCs as covariates for all our genetic analyses.

The number of loci were counted from the GWAS results through a chromosome-wide clumping procedure. The top significantly detected SNP from one chromosome was tallied as a hit, then all significant hits within 1 MB from the SNP were ignored. The procedure was repeated for any remaining significant detection in the chromosome, then repeated within all chromosomes.

### GWAS on AutoComplete-imputed phenotypes

For the imputation of phenotypes for which we performed GWAS, AutoComplete was allowed to fit all available individuals in order to impute missing entries. For binary phenotypes, phenotypes were imputed in a continuous range of 0 ~ 1 reflective of confidence in the prediction.

GWAS on directly-phenotyped and imputed phenotypes in UKBB was performed using imputed genotype data at the 5,776,313 SNPs (minor allele frequency > 5%, INFO score > 0.9) using logistic regression and linear regression implemented in PLINK v2 [53] for binary and quantitative phenotypes respectively. For imputed phenotypes, linear regression was performed.

### Additional analysis of imputed phenotypes

The effective sample size was calculated as a function of imputation accuracy for a given phenotype from simulations (1% missingness) and the number of missing values imputed, such that 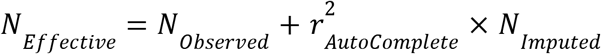 for a given phenotype.

We examined genetic correlations (*r_G_*) between a subset of phenotypes within the psychiatric disorder dataset collected within the UK Biobank, and related phenotypes collected from cohorts outside the UK Biobank. The five phenotypes examined within the UK Biobank were: LifetimeMDD [51], anxiety with drug use, using medication for MDD, number of MDD episodes, and having ever taken cannabis. In the context of these phenotypes, we gathered GWAS summary statistics from external studies which examined cannabis-use disorder [14], multiple psychiatric disorders [15], MDD [16], schizophrenia [17], and bipolar disorder [18]. We used LD Score regression (LDSC) [19] to estimate *r_G_* between each pairing of phenotypes using LD Scores estimated from the 1000 Genomes white European population [26,27].

## Data Availability

The genotype and phenotype data are available by application from the UKBB https://www.ukbiobank.ac.uk. The LD Scores from the 1000 Genomes project are available from https://alkesgroup.broadinstitute.org/LDSCORE/.

## URLs

AutoComplete software: https://github.com/sriramlab/AutoComplete

Plink 2.0: https://www.cog-genomics.org/plink/2.0/

LDSC: https://github.com/bulik/ldsc

HI-VAE: https://github.com/probabilistic-learning/HI-VAE

GAIN: https://github.com/jsyoon0823/GAIN

kNN: https://scikit-learn.org/stable/modules/generated/sklearn.impute.KNNImputer.html

MissForeset: https://cran.r-project.org/web/packages/missForest/index.html

MICE: https://cran.r-project.org/web/packages/mice/index.html

## Acknowledgments

This research was conducted using the UK Biobank Resource under applications 33127 and 33297. We thank the participants of UK Biobank for making this work possible. This work was funded by NIH grants R01HG010505 and R01DK132775 (P.P., M.A.), R35GM125055 (S.S.), and NSF grants III-1705121 (S.S., U.A). This work was also funded by Lundbeckfonden Fellowship R335-2019-2318.

## References

1. Greenland S, Finkle WD. A Critical Look at Methods for Handling Missing Covariates in Epidemiologic Regression Analyses. American Journal of Epidemiology [Internet]. 1995 Dec;142(12):1255–64. Available from: https://doi.org/10.1093/oxfordjournals.aje.a117592

2. Rubin DB. Multiple imputation for nonresponse in surveys [Internet]. Wiley; 2004. (Wiley classics library). Available from: https://books.google.com/books?id=bQBtw6rx\_mUC

3. Goodfellow I, Pouget-Abadie J, Mirza M, Xu B, Warde-Farley D, Ozair S, et al. Generative adversarial nets. In: Ghahramani Z, Welling M, Cortes C, Lawrence N, Weinberger KQ, editors. Advances in neural information processing systems [Internet]. Curran Associates, Inc.; 2014. Available from: https://proceedings.neurips.cc/paper/2014/file/5ca3e9b122f61f8f06494c97b1afccf3-Paper.pdf

4. Kingma DP, Welling M. An introduction to variational autoencoders. Foundations and Trends^®^ in Machine Learning [Internet]. 2019;12(4):307–92. Available from: http://dx.doi.org/10.1561/2200000056

5. Bycroft C, Freeman C, Petkova D, Band G, Elliott LT, Sharp K, et al. The UK biobank resource with deep phenotyping and genomic data. Nature [Internet]. 2018 Oct 1;562(7726):203–9. Available from: https://doi.org/10.1038/s41586-018-0579-z

6. Stekhoven DJ, Bühlmann P. MissForest—non-parametric missing value imputation for mixed-type data. Bioinformatics [Internet]. 2011 Oct;28(1):112–8. Available from: https://doi.org/10.1093/bioinformatics/btr597

7. Buuren S van. Flexible imputation of missing data. Second edition. Boca Raton, FL.: CRC Press; 2018.

8. Troyanskaya O, Cantor M, Sherlock G, Brown P, Hastie T, Tibshirani R, et al. Missing value estimation methods for DNA microarrays. Bioinformatics [Internet]. 2001 Jun;17(6):520–5. Available from: https://doi.org/10.1093/bioinformatics/17.6.520

9. Hastie T, Mazumder R, Lee JD, Zadeh R. Matrix completion and low-rank SVD via fast alternating least squares. Journal of machine learning research: JMLR [Internet]. 2015;16:3367–402. Available from: https://pubmed.ncbi.nlm.nih.gov/31130828

10. Yoon J, Jordon J, Schaar M van der. GAIN: Missing data imputation using generative adversarial nets. In: Dy J, Krause A, editors. Proceedings of the 35th international conference on machine learning [Internet]. PMLR; 2018. p. 5689–98. (Proceedings of machine learning research; vol. 80). Available from: https://proceedings.mlr.press/v80/yoon18a.html

11. Nazábal A, Olmos PM, Ghahramani Z, Valera I. Handling incomplete heterogeneous data using VAEs. Pattern Recognition [Internet]. 2020;107:107501. Available from: https://www.sciencedirect.com/science/article/pii/S0031320320303046

12. Pritchard JK, Przeworski M. Linkage disequilibrium in humans: Models and data. The American Journal of Human Genetics. 2001;69(1):1–4.

13. Littlejohns TJ, Holliday J, Gibson LM, Garratt S, Oesingmann N, Alfaro-Almagro F, et al. The UK biobank imaging enhancement of 100,000 participants: Rationale, data collection, management and future directions. Nat Commun. 2020 May;11(1):2624.

14. Johnson EC, Demontis D, Thorgeirsson TE, Walters RK, Polimanti R, Hatoum AS, et al. A large-scale genome-wide association study meta-analysis of cannabis use disorder. Lancet Psychiatry. 2020 Oct;7(12):1032–45.

15. Cross-Disorder Group of the Psychiatric Genomics Consortium. Identification of risk loci with shared effects on five major psychiatric disorders: A genome-wide analysis. Lancet. 2013 Feb;381(9875):1371–9.

16. Major Depressive Disorder Working Group of the Psychiatric GWAS Consortium, Ripke S, Wray NR, Lewis CM, Hamilton SP, Weissman MM, et al. A mega-analysis of genome-wide association studies for major depressive disorder. Mol Psychiatry. 2012 Apr;18(4):497–511.

17. Schizophrenia Working Group of the Psychiatric Genomics Consortium. Biological insights from 108 schizophrenia-associated genetic loci. Nature. 2014 Jul;511(7510):421–7.

18. Psychiatric GWAS Consortium Bipolar Disorder Working Group. Large-scale genome-wide association analysis of bipolar disorder identifies a new susceptibility locus near ODZ4. Nat Genet. 2011 Sep;43(10):977–83.

19. Bulik-Sullivan BK, Loh PR, Finucane HK, Ripke S, Yang J, Patterson N, et al. LD score regression distinguishes confounding from polygenicity in genome-wide association studies. Nature Genetics [Internet]. 2015 Mar 1;47(3):291–5. Available from: https://doi.org/10.1038/ng.3211

20. Bulik-Sullivan B, Finucane HK, Anttila V, Gusev A, Day FR, Loh PR, et al. An atlas of genetic correlations across human diseases and traits. Nature Genetics [Internet]. 2015 Nov 1;47(11):1236–41. Available from: https://doi.org/10.1038/ng.3406

21. He K, Zhang X, Ren S, Sun J. Deep residual learning for image recognition. In: Proceedings of the IEEE conference on computer vision and pattern recognition. 2016. p. 770–8.

22. Ding Z, Xu Y, Xu W, Parmar G, Yang Y, Welling M, et al. Guided variational autoencoder for disentanglement learning. In: Proceedings of the IEEE/CVF conference on computer vision and pattern recognition. 2020. p. 7920–9.

23. Lundberg SM, Lee SI. A unified approach to interpreting model predictions. In: Guyon I, Luxburg UV, Bengio S, Wallach H, Fergus R, Vishwanathan S, et al., editors. Advances in neural information processing systems 30 [Internet]. Curran Associates, Inc.; 2017. p. 4765–74. Available from: http://papers.nips.cc/paper/7062-a-unified-approach-to-interpreting-model-predictions.pdf

24. Wilman HR, Kelly M, Garratt S, Matthews PM, Milanesi M, Herlihy A, et al. Characterisation of liver fat in the UK biobank cohort. PLoS One. 2017 Feb;12(2):e0172921.

25. Zhou P, Feng J, Ma C, Xiong C, Hoi SCH, E W. Towards theoretically understanding why sgd generalizes better than adam in deep learning. In: Larochelle H, Ranzato M, Hadsell R, Balcan MF, Lin HT, editors. Advances in neural information processing systems 33: Annual conference on neural information processing systems 2020, NeurIPS 2020, december 6-12, 2020, virtual [Internet]. 2020. Available from: https://proceedings.neurips.cc/paper/2020/hash/f3f27a324736617f20abbf2ffd806f6d-Abstract.html

26. Auton A, Abecasis GR, Altshuler DM, Durbin RM, Bentley DR, Chakravarti A, et al. A global reference for human genetic variation. Nature [Internet]. 2015 Oct 1;526(7571):68–74. Available from: https://doi.org/10.1038/nature15393

27. Zheng J, Erzurumluoglu AM, Elsworth BL, Kemp JP, Howe L, Haycock PC, et al. LD Hub: a centralized database and web interface to perform LD score regression that maximizes the potential of summary level GWAS data for SNP heritability and genetic correlation analysis. Bioinformatics [Internet]. 2016 Sep;33(2):272–9. Available from: https://doi.org/10.1093/bioinformatics/btw613

28. Beretta L, Santaniello A. Nearest neighbor imputation algorithms: A critical evaluation. BMC Medical Informatics and Decision Making [Internet]. 2016 Jul 25;16(3):74. Available from: https://doi.org/10.1186/s12911-016-0318-z

29. Little RJ, Rubin DB. Statistical analysis with missing data. Vol. 793. John Wiley & Sons; 2019.

30. Städler N, Bühlmann P. Missing values: Sparse inverse covariance estimation and an extension to sparse regression. Statistics and Computing [Internet]. 2012 Jan;22(1):219–35. Available from: https://doi.org/10.1007/s11222-010-9219-7

31. Städler N, Stekhoven DJ, Bühlmann P. Pattern alternating maximization algorithm for missing data in high-dimensional problems. J Mach Learn Res. 2014 Jan;15(1):1903–28.

32. Allen GI, Tibshirani R. Transposable regularized covariance models with an application to missing data imputation. The Annals of Applied Statistics [Internet]. 2010;4(2):764–90. Available from: https://doi.org/10.1214/09-AOAS314

33. Mongia A, Sengupta D, Majumdar A. McImpute: Matrix completion based imputation for single cell RNA-seq data. Frontiers in Genetics [Internet]. 2019;10:9. Available from: https://www.frontiersin.org/article/10.3389/fgene.2019.00009

34. Phung S, Kumar A, Kim J. A deep learning technique for imputing missing healthcare data. In: 2019 41st annual international conference of the IEEE engineering in medicine and biology society (EMBC). 2019. p. 6513–6.

35. Koren Y, Bell R, Volinsky C. Matrix factorization techniques for recommender systems. Computer. 2009;42(8):30–7.

36. Mazumder R, Hastie T, Tibshirani R. Spectral regularization algorithms for learning large incomplete matrices. Journal of Machine Learning Research [Internet]. 2010;11(80):2287–322. Available from: http://jmlr.org/papers/v11/mazumder10a.html

37. Dahl A, Iotchkova V, Baud A, Johansson Å, Gyllensten U, Soranzo N, et al. A multiple-phenotype imputation method for genetic studies. Nature Genetics [Internet]. 2016 Apr 1;48(4):466–72. Available from: https://doi.org/10.1038/ng.3513

38. Qiu YL, Zheng H, Gevaert O. Genomic data imputation with variational auto-encoders. GigaScience [Internet]. 2020 Aug;9(8). Available from: https://doi.org/10.1093/gigascience/giaa082

39. García-Laencina PJ, Sancho-Gómez JL, Figueiras-Vidal AR. Pattern classification with missing data: A review. Neural Computing and Applications [Internet]. 2010 Mar 1;19(2):263–82. Available from: https://doi.org/10.1007/s00521-009-0295-6

40. Srivastava A, Valkov L, Russell C, Gutmann MU, Sutton C. VEEGAN: Reducing mode collapse in GANs using implicit variational learning. In: Guyon I, Luxburg UV, Bengio S, Wallach H, Fergus R, Vishwanathan S, et al., editors. Advances in neural information processing systems [Internet]. Curran Associates, Inc.; 2017. Available from: https://proceedings.neurips.cc/paper/2017/file/44a2e0804995faf8d2e3b084a1e2db1d-Paper.pdf

41. Beaulieu-Jones BK, Moore JH. MISSING DATA IMPUTATION IN THE ELECTRONIC HEALTH RECORD USING DEEPLY LEARNED AUTOENCODERS. Pacific Symposium on Biocomputing Pacific Symposium on Biocomputing [Internet]. 2017;22:207–18. Available from: https://doi.org/10.1142/9789813207813_0021

42. Kollewe K, Mauss U, Krampfl K, Petri S, Dengler R, Mohammadi B. ALSFRS-r score and its ratio: A useful predictor for ALS-progression. Journal of the Neurological Sciences [Internet]. 2008 Dec 15;275(1):69–73. Available from: https://doi.org/10.1016/j.jns.2008.07.016

43. Arisdakessian C, Poirion O, Yunits B, Zhu X, Garmire LX. DeepImpute: An accurate, fast, and scalable deep neural network method to impute single-cell RNA-seq data. Genome Biology [Internet]. 2019 Oct 18;20(1):211. Available from: https://doi.org/10.1186/s13059-019-1837-6

44. Vincent P, Larochelle H, Bengio Y, Manzagol PA. Extracting and composing robust features with denoising autoencoders. In: Proceedings of the 25th international conference on machine learning [Internet]. New York, NY, USA: Association for Computing Machinery; 2008. p. 1096–103. (ICML ‘08). Available from: https://doi.org/10.1145/1390156.1390294

45. Vincent P, Larochelle H, Lajoie I, Bengio Y, Manzagol PA. Stacked denoising autoencoders: Learning useful representations in a deep network with a local denoising criterion. Journal of Machine Learning Research [Internet]. 2010;11(110):3371–408. Available from: http://jmlr.org/papers/v11/vincent10a.html

46. Miotto R, Li L, Kidd BA, Dudley JT. Deep patient: An unsupervised representation to predict the future of patients from the electronic health records. Scientific Reports [Internet]. 2016 May 17;6(1):26094. Available from: https://doi.org/10.1038/srep26094

47. Wu R, Zhang A, Ilyas I, Rekatsinas T. Attention-based learning for missing data imputation in HoloClean. In: Dhillon I, Papailiopoulos D, Sze V, editors. Proceedings of machine learning and systems [Internet]. 2020. p. 307–25. Available from: https://proceedings.mlsys.org/paper/2020/file/202cb962ac59075b964b07152d234b70-Paper.pdf

48. Spinelli I, Scardapane S, Uncini A. Missing data imputation with adversarially-trained graph convolutional networks. Neural Networks [Internet]. 2020;129:249–60. Available from: https://www.sciencedirect.com/science/article/pii/S0893608020302185

49. Kyono T, Zhang Y, Bellot A, Van der Schaar M. MIRACLE: Causally-aware imputation via learning missing data mechanisms. In: Advances in neural information processing systems [Internet]. Curran Associates, Inc.; 2021. Available from: https://proceedings.neurips.cc//paper/2021/hash/c80bcf42c220b8f5c41f85344242f1b0-Abstract.html

50. Hormozdiari, Farhad, Eun Yong Kang, Michael Bilow, Eyal Ben-David, Chris Vulpe, Stela McLachlan, Aldons J. Lusis, Buhm Han, and Eleazar Eskin. “Imputing phenotypes for genome-wide association studies.” The American Journal of Human Genetics 99, no. 1 (2016): 89–103.

51. Cai, Na, Joana A. Revez, Mark J. Adams, Till FM Andlauer, Gerome Breen, Enda M. Byrne, Toni-Kim Clarke et al. “Minimal phenotyping yields genome-wide association signals of low specificity for major depression.” Nature Genetics 52, no. 4 (2020): 437–447.

52. Abraham, Gad, Yixuan Qiu, and Michael Inouye. “FlashPCA2: principal component analysis of Biobank-scale genotype datasets.” Bioinformatics (2017).

53. Chang, Christopher C., Carson C. Chow, Laurent CAM Tellier, Shashaank Vattikuti, Shaun M. Purcell, and James J. Lee. “Second-generation PLINK: rising to the challenge of larger and richer datasets.” Gigascience 4, no. 1 (2015): s13742–015.

54. Sundararajan, Mukund, Ankur Taly, and Qiqi Yan. “Axiomatic attribution for deep networks.” In International conference on machine learning, pp. 3319–3328. PMLR, 2017.

55. Williams, Valerie F., Stephen B. Taubman, and Shauna Stahlman. “Non-alcoholic fatty liver disease (NAFLD), active component, US Armed Forces, 2000-2017.” MSMR 26, no. 1 (2019): 2–11.

56. Miao, Zong, Kristina M. Garske, David Z. Pan, Amogha Koka, Dorota Kaminska, Ville Männistö, Janet S. Sinsheimer, Jussi Pihlajamäki, and Päivi Pajukanta. “Identification of 90 NAFLD GWAS loci and establishment of NAFLD PRS and causal role of NAFLD in coronary artery disease.” Human Genetics and Genomics Advances 3, no. 1 (2022): 100056.

57. Liang, Z., Zhang, G., Huang, J. X. & Hu, Q. V. Deep learning for healthcare decision making with EMRs. IEEE BIBM, 556–559 (2014).

58. Bengio, Y., Courville, A. & Vincent, P. Representation learning: a review and new perspectives. IEEE T. Pattern Anal. Mach. Intell. 35, 1798–1828 (2013).

59. LeCun, Y., Bengio, Y. & Hinton, G. Deep learning. Nature 521, 436–444 (2015).

60. Helmstaedter, M. et al. Connectomic reconstruction of the inner plexiform layer in the mouse retina. Nature 500, 168–174 (2013).

61. Leung, M. K. K., Xiong, H. Y., Lee, L. J. & Frey, B. J. Deep learning of the tissue-regulated splicing code. Bioinformatics 30, 121–129 (2014).

62. Xiong, H. Y. et al. The human splicing code reveals new insights into the genetic determinants of disease. Science 347, 144–151 (2015).

63. Alipanahi, B., Delong, A., Weirauch, M. T. & Frey, B. J. Predicting the sequence specificities of DNA-and RNA-binding proteins by deep learning. Nature Biotech. 33, 831–838 (2015).

64. Ma, J. S., Sheridan, R. P., Liaw, A., Dahl, G. E. & Svetnik, V. Deep neural nets as a method for quantitative structure-activity relationships. J. Chem. Inf. Model 55, 263–274 (2015).

65. Zeggini E, Scott LJ, Saxena R, Voight BF, Marchini JL, et al. (2008) Meta-analysis of genome-wide association data and large-scale replication identifies additional susceptibility loci for type 2 diabetes. Nat Genet 40: 638–645.

66. Marchini J, Howie B, Myers S, McVean G, Donnelly P (2007) A new multipoint method for genome-wide association studies by imputation of genotypes. Nat Genet 39: 906–913.

67. Khera, Amit V., Mark Chaffin, Krishna G. Aragam, Mary E. Haas, Carolina Roselli, Seung Hoan Choi, Pradeep Natarajan et al. “Genome-wide polygenic scores for common diseases identify individuals with risk equivalent to monogenic mutations.” Nature genetics 50, no. 9 (2018): 1219–1224.

68. Gigantesco, Antonella, and Pierluigi Morosini. “Development, reliability and factor analysis of a self-administered questionnaire which originates from the World Health Organization’s Composite International Diagnostic Interview–Short Form (CIDI-SF) for assessing mental disorders.” Clinical Practice and Epidemiology in Mental Health 4, no. 1 (2008): 1–10.

69. Kroenke, Kurt, and Robert L. Spitzer. “The PHQ-9: a new depression diagnostic and severity measure.” Psychiatric annals 32, no. 9 (2002): 509–515.

70. Li, Ting, Zheng Ning, and Xia Shen. “Improved estimation of phenotypic correlations using summary association statistics.” Frontiers in genetics (2021): 1291.

71. Van Buuren, Stef, and Karin Oudshoorn. Flexible multivariate imputation by MICE. Leiden: TNO, 1999.

