## Supplementary Material for "Deep Learning-based Phenotype Imputation on Population-scale Biobank Data Increases Genetic Discoveries"

### Supplementary Material: Deep Learning-based Multiple Phenotype Imputation on Biobank-scale Data Increases Genetic Discoveries

#### S1 Related Work

When a subset of features are missing consistently, the missing values can be predicted using the remaining observed features using standard supervised learning. In practice, patterns of missingness can be complex. Multivariate imputation by chained equations (MICE) [71], which aims to repeatedly fit a conditional distribution for each feature given the other and using this distribution to impute missing values, has emerged as a principled framework to dealing with missing data. Several variants of this approach, based on how the conditional distributions are modeled, have been developed and methods which leverage random forests within MICE such as MissForest [6] and MICE-Forest [7] are widely used. These approaches have the advantage of being able to handle mixed data types. K-Nearest Neighbors [8] is also a prevalent approach for imputation of continuous data (originally introduced to impute gene expression data). This approach attempts to impute a missing phenotype based on K "nearest" phenotypes (determined by computing a distance on the observed phenotypes). The accuracy of this approach depends on the choice of K and appropriate choice of a measure of distance across phenotypes [28].

Approaches such as the Multivariate Normal Model [29], MissGLasso [30], MissPALasso [31], and TRCMA [32] aim to estimate the parameters of the distribution of the data, in some cases with additional regularization. MissGLImp and MissPALasso aim to learn a sparse inverse-covariance matrix underlying the partially observed phenotypes using a EM-style algorithm while TRCMA aims to fit a matrix-normal model to the matrix of phenotypes over individuals (with some entries missing) where the covariance matrices underlying the matrix normal model are regularized. SoftImpute [9] has become one of the most prevalent imputation methods due to its scalability and flexibility [33,34]. The method builds upon work in matrix completion, including SVD-based imputation [35] and HardImpute [36]. It assumes that the latent phenotype matrix has low-rank that it attempts to estimate by searching for a matrix that is close to the observed entries of the phenotype matrix while also having an approximately low rank (as quantified by its nuclear norm).

In the context of genetic studies, PHENIX [37] models the matrix of phenotypes observed across individuals (with some entries missing) as arising from a matrix-normal distribution. This elegant approach accounts for the genetic relatedness (that can induce correlations for a given phenotype measured across individuals) and pleiotropy (that can induce correlations across phenotypes measured in the same individual). The parameters of the model are estimated using a variational Bayes algorithm which also provides an approximate posterior distribution over the missing phenotypes. While an elegant model, it is challenging to scale PHENIX to large numbers of individuals and phenotypes (for example, this would require eigendecomposition of the kinship or genetic relatedness matrix). Further, the model underlying PHENIX is designed for normally-distributed phenotypes (the method was shown to be less accurate on non-normally distributed phenotypes though it remained the most accurate compared to other methods) [37].

PhenIMP [50] considers the setting where phenotypes that are closely related to a phenotype of interest have been collected in large samples. PhenIMP models the joint distribution of the phenotype of interest and related phenotypes in each individual as arising independently from a zero-mean multivariate normal distribution. The covariance matrix of the distribution over the phenotypes is estimated from a dataset where all phenotypes are measured which can then be used to impute the phenotype of interest in datasets where only the related phenotypes are measured. PhenIMP requires access to a complete dataset on which the phenotype of interest and related phenotypes are measured and is also designed for normally distributed phenotypes.

Recent advancements in deep-learning have given rise to deep generative models which are capable of learning high-dimensional, multi-modal distributions. Notably, Variational Auto-Encoders (VAE) offer one framework of using Deep Neural Nets (DNN) to build a probabilistic model [4]. Variants of VAE have been proposed for the imputation problem [38] though the proposed approach assumes the presence of samples with complete observations to learn the model parameters. HI-VAE [11] defines a comprehensive deep probabilistic model which accounts for missingness in observations, as generated from a mixture distribution, and natively supports various heterogeneous data formats (continuous, binary, categorical, ordinal). The flexibility of the model makes it one of the most suitable deep-learning methods in the context of medical data which may consist of all the noted attributes. In addition to VAEs, Generative Adversarial Networks (GAN) [3] offer an alternative approach to generative modeling using a deep generator-discriminator paired architecture. GAIN [10] extends GANs to the imputation problem and has been shown to obtain improved accuracy over MICE and missForest [6,7,39]. However HI-VAE has been shown to be favorable in comparison to GAIN under varying datasets [11] and the lack of consistency in the convergence of GANs has been one barrier to their broader use [40].

Among deep-learning methods, Auto Encoders (AE) have remained a competitive approach for imputation. The method is based on discriminative training of deep encoder-decoder neural nets with a focus on reconstructing perturbed or missing values. Several works have used AEs for imputing medical records. The earlier works, however, relied on mostly complete datasets with high proportion of observed values, such that underlying missingness was not a consideration for the methods, and their utility was mostly demonstrated for synthetically created levels of missing data. *Beaulieu-Jones et al.* [41] demonstrated that medical records relating to ALS [42] could be imputed with higher accuracy using AE than several non deep-learning methods. However, the work was defined specifically for one data format (binary labels), and evaluated on a synthetic dataset of missing values generated from 2000 individuals. DeepImpute [43] similarly demonstrated the applicability of AEs in on single-cell RNA-seq data where completely observed samples were available, evaluating on incomplete datasets which were simulated.

*Phung et al.* [34] proposed a denoising auto-encoder [44,45] where Gaussian noise was dynamically added to the data, successfully improving imputation for infant mortality records. DeepPatient [46] leveraged stacked DAEs under uniform masking noise to fit medical records. While not explicitly defined for imputation, this work demonstrated that the latent representations learned by DAEs were predictive of various diseases.

Novel deep neural-net architectures are continually developed such as attention-based [47], graph-based [48], or causally regularized [49] methods, but several aspects of AutoEncoders, the effectiveness of denoising, and their extension to highly missing, massive datasets for real-world impact have yet to be explored. Our work emphasizes the strong performance and reliability of DAEs, and we arrive at an imputing DAE which is favorable to many conventional and deep-learning approaches which generalizes to the types of missingness found in biobank-scale data.

#### S2 Tests in Smaller Scales

We organized a smaller subset of 86 phenotypes related to blood lab measurements within the cardiometabolic dataset called the Blood Labs dataset to test a wider variety of methods which could not scale to the size of the two main datasets with hundreds of phenotypes. This dataset contained 68 continuous and 18 binary phenotypes, with the phenotype with highest missingness being 91% missing. There were 291,273 individuals in total for this dataset, where 151,273 individuals were randomly selected as the training set and the remaining individuals were reserved for testing. A smaller subset was formed from the initial split by drawing 5,000 individuals each from the initial training and testing splits. In this setting, we could perform fitting and hyperparameter search for two widely applied methods MissForest [6] and MICE Forest [2] in reasonable time. MissForest was configured to use up to 10 trees and MICE Forest was configured to use up to 100 trees.

In evaluating the smaller subset, a similar trend in accuracy was observed in comparison to the full dataset for all methods (Figure S5). Given reduced sample sizes, the confidence intervals also grew larger. In the smaller setting, AutoComplete and SoftImpute appeared the top two most favorable methods in imputation accuracy. We observed that AutoComplete-UKBB which had learned from all the data obtained roughly a higher accuracy (point estimate), an increase from 0.324 to 0.360 from the top two methods (+11%) at 1% simulated missingness. Between AutoComplete and MICE Forest, an improvement from 0.278 to 0.324 (+16%) was observed while MissForest obtained an accuracy of 0.224 (+47%). MissForest and MICE Forest appeared much more favorable than K-Nearest Neighbors and the generative-adversarial method GAIN. The trends persisted for additional simulated settings of missingness up to 50%.

#### S3 Contribution of copy-masking

We performed ablation tests to determine that copy-masking is a key factor in improving imputation accuracy for AutoComplete. We measured overall performance using average $r^2$ across phenotypes from the psychiatric disorders dataset with increasing percentages of simulated missingness (1% ∼ 50% missing). We compared our method with training the denoising architecture with uniform random masking of observed values in increasing amounts of 10% ∼ 90% (Figure S6). For the simulated setting of 1% missing values, the highest average $r^2$ obtained through uniform masking was 0.121 in comparison to 0.142 with AutoComplete (17%) with similar trends in tests with increasing missingness (average 16% improvement). We therefore conclude that AutoComplete benefits substantially from being trained on realistic missingness patterns that aid the denoising behavior of the deep learning model.

We tested the effect of not using copy-masking in terms of evaluating the imputation methods. We simulated missingness for testing using uniform random missingness in the range of 1% to 50% of the total observed measurements. We evaluated the same AutoComplete and SoftImpute models from our main experiments in this setting (Figure S7). We observe that copy-masking led to a substantially higher imputation accuracy for the psychiatric disorders dataset (0.213 in $r^2$) in comparison to using copy-masking (0.116). Examining a highly missing phenotype LifetimeMDD of clinical interest, imputation accuracy was close to 100% for 1% missingness simulation for AutoComplete and SoftImpute (0.987 and 0.926), which was highly inflated in comparison to our evaluations with copy-masking (0.468 for AutoComplete and 0.311 for SoftImpute). While imputation accuracies were overall inflated for this setting, AutoComplete imputations were more accurate than SoftImpute imputations on average (0.213 and 0.201 respectively).

### Supplementary Tables and Figures

| Dataset | N | # Pheno. | # Cont. | # Binary | $\varnothing_N$ | $\varnothing_P$ |
|---|---|---|---|---|---|---|
| Cardiometabolic | 285,405 | 230 (49%) | 46 (7%) | 184 (60%) | 50% (78%) | 47% (99%) |
| Psychiatric Disorders | 337,126 | 372 (46%) | 98 (55%) | 274 (42%) | 53% (77%) | 67% (99%) |

**Table S1:** We collected two sets of phenotypes from two studies related to the UK Biobank. Each dataset contains hundreds of thousands of individuals (*N*) and a heterogeneous mix of continuous (*Cont.*) and binary valued phenotypes. Percent of all values which are missing for select phenotypes are reported in parentheses after the number (#) of such traits. We report the median percentage of measurements missing per individual ($Ø_N$) (maximum missing in parentheses) in addition to the median percentage of measurements missing per phenotype ($Ø_P$).

| | | Cardiometabolic | | | | | Psychiatric Disorders | | | | |
|---|---|---|---|---|---|---|---|---|---|---|---|
| | Method | 1% | 5% | 10% | 20% | 50% | 1% | 5% | 10% | 20% | 50% |
| $r^2$ | SI | 6 (0) | 31 (7) | 35 (9) | 23 (11) | 70 (11) | 4 (0) | 21 (2) | 71 (2) | 129 (2) | 148 (31) |
| | HIVAE | 11 (0) | 86 (0) | 131 (1) | 151 (2) | - (-) | 20 (0) | 63 (0) | 97 (0) | 143 (0) | - (-) |
| | GAIN | 102 (0) | 156 (0) | 169 (0) | 177 (0) | 177 (0) | 120 (0) | 174 (0) | 215 (0) | 235 (1) | - (-) |
| | KNN | 85 (0) | 148 (0) | 154 (0) | 163 (0) | 162 (0) | 108 (0) | 173 (0) | 203 (0) | 236 (0) | 223 (1) |
| AUPR | SI | 0 (0) | 3 (0) | 3 (0) | 5 (0) | 4 (0) | 0 (0) | 3 (0) | 21 (0) | 61 (0) | 62 (24) |
| | HIVAE | 1 (0) | 4 (0) | 4 (0) | 4 (0) | - (-) | 2 (0) | 16 (0) | 28 (0) | 59 (0) | - (-) |
| | GAIN | 4 (0) | 6 (0) | 8 (0) | 11 (0) | 13 (0) | 54 (0) | 98 (2) | 126 (5) | 141 (6) | - (-) |
| | KNN | 4 (0) | 5 (0) | 8 (0) | 7 (0) | 11 (0) | 34 (2) | 83 (2) | 102 (10) | 114 (18) | 68 (71) |
| AUROC | SI | 2 (0) | 3 (0) | 5 (0) | 6 (0) | 6 (0) | 3 (0) | 17 (1) | 58 (0) | 81 (0) | 86 (30) |
| | HIVAE | 2 (0) | 2 (0) | 5 (0) | 3 (0) | - (-) | 8 (1) | 31 (0) | 43 (1) | 76 (0) | - (-) |
| | GAIN | 5 (0) | 9 (0) | 9 (0) | 12 (0) | 13 (0) | 91 (0) | 127 (0) | 155 (0) | 167 (0) | - (-) |
| | KNN | 5 (0) | 8 (0) | 11 (0) | 13 (0) | 13 (0) | 92 (1) | 143 (0) | 157 (0) | 179 (0) | 179 (0) |

**Table S2:** Number of phenotypes for which AutoComplete significantly improved imputation accuracy relative to other methods. Significant decreases in accuracy are indicated in parentheses. Imputation accuracy was determined through a simulation of 1%~50% missingness on the cardiometabolic and psychiatric disorders datasets, for the metrics of squared Pearson correlation ($r^2$), Area Under Precision Recall (AUPR), and Area Under Receiver Operating Characteristic (AUROC). A subset of methods could not be reliability fit and compared for high degrees of missingness (-).

| Phenotype | N | $r^2$ | N Imputed | N Effective | Increase |
|---|---|---|---|---|---|
| LifetimeMDD | 67,164 | 0.47 | 269,962 | 193,379 | 187% |
| Anxiety with Drug Use | 18,665 | 0.19 | 318,461 | 79,16 | 323% |
| Medication for MDD | 32,267 | 0.35 | 304,859 | 139,167 | 331% |
| MDD Episodes | 58,994 | 0.18 | 278,132 | 110,518 | 87% |
| Cannabis Ever Taken | 110,188 | 0.24 | 226,938 | 165,291 | 50% |

**Table S3:** Increase in effective sample sizes from AutoComplete imputations of highly missing phenotypes in the psychiatric disorders dataset. Of 337,126 total individuals in the psychiatric disorders dataset, the available sample size of each phenotype (N) was much smaller due to missingness. All missing measurements were imputed ($N_{Imputed}$) using AutoComplete. The accuracy of the imputation method for each phenotype could be measured in simulations ($r^2$), allowing an approximation of the hypothetical effective sample size ($N_{Effective} = N + r^2 \times N_{Imputed}$) and an estimate of the increase in sample size (Increase).

(a)

| Method | CPU | GPU | H | 10% | 25% | 50% | 100% |
|---|---|---|---|---|---|---|---|
| AutoComplete | · | ✓ | ✓ | 41s | 103s | 387s | 1190s |
| KNN [8] | · | · | ✓ | 45s | 393s | 12040s | · |
| SoftImpute [36] | · | · | . | 64s | 380s | 837s | 1659s |
| GAIN [10] | · | ✓ | . | 4.2s | 12s | 27s | 87s |
| HI-VAE [11] | · | ✓ | ✓ | 46s | 272s | 1028s | 5000s |
| missForest [6] | ✓ | · | ✓ | 192s | 4998s | · | · |
| MICE [7] | ✓ | · | ✓ | 292s | 3703s | 31373s | · |

(b)

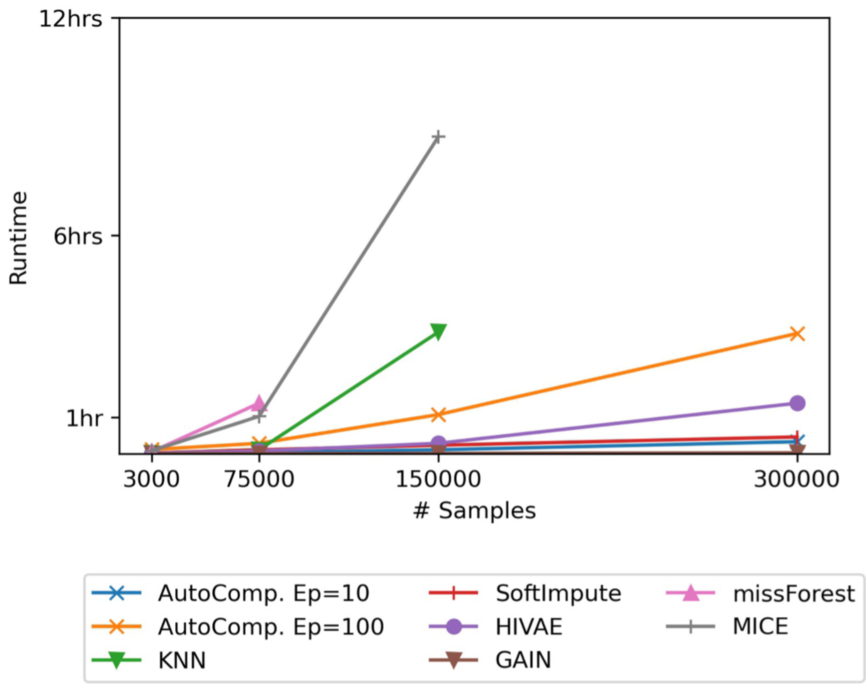


**Figure S1:** (a) Table showing the time needed to fit each method (either 10 iterations or until a default convergence criteria). Times were measured for increasing fractions of the Psychiatric disorders dataset (10%~100%). The availability of either CPU or GPU acceleration is reported for each method. In addition methods which natively support mixed data types are indicated (H). Benchmarks were terminated ("·") if more than 12 hrs elapsed in fitting the method. (b) The time to fit each method is illustrated, including the training of AutoComplete for 10 epochs and up to 100 epochs.

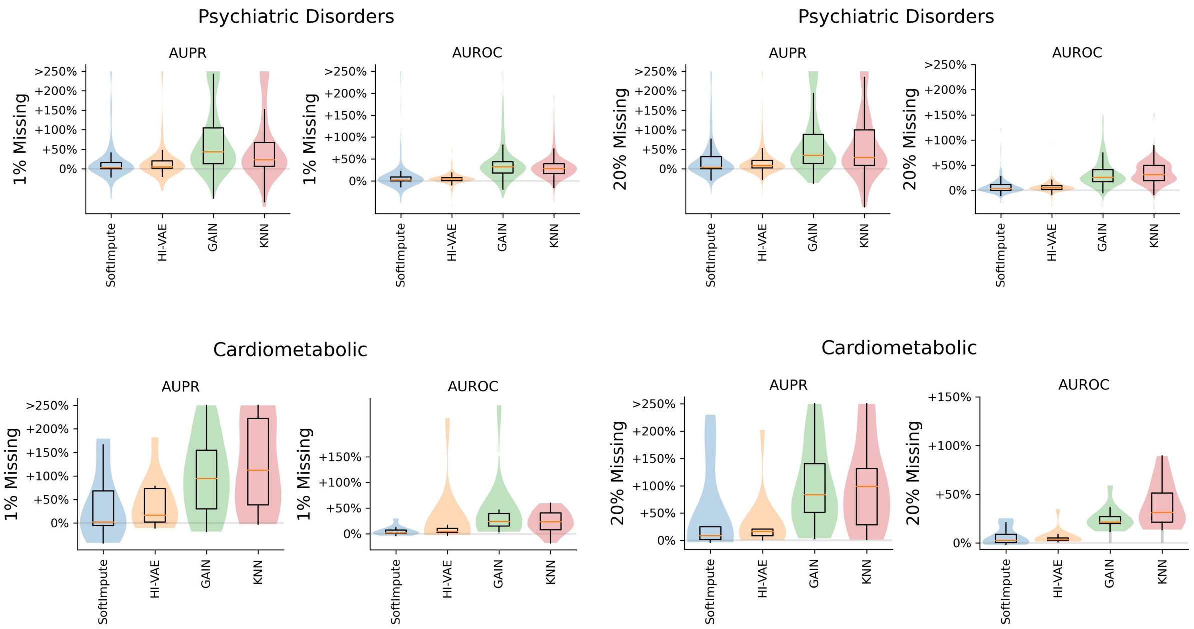


**Figure S2:** Relative gains in accuracy of AutoComplete for binary phenotypes under the metrics: Area Under Precision Recall Curve (AUPR), and Area Under Receiver Operating Characteristic Curve (AUROC). Comparisons were made to SoftImpute, HI-VAE, GAIN and KNN, highlighted for the case of 1% and 20% simulated missingness of the original data. Relative change in accuracy was determined as e.g. $\Delta = (\theta_{AutoComplete} - \theta_{SoftImpute})/\theta_{SoftImpute}$ for score θ. GAIN could not be reliability trained for 20% simulated missingness and beyond for the psychiatric disorders dataset.

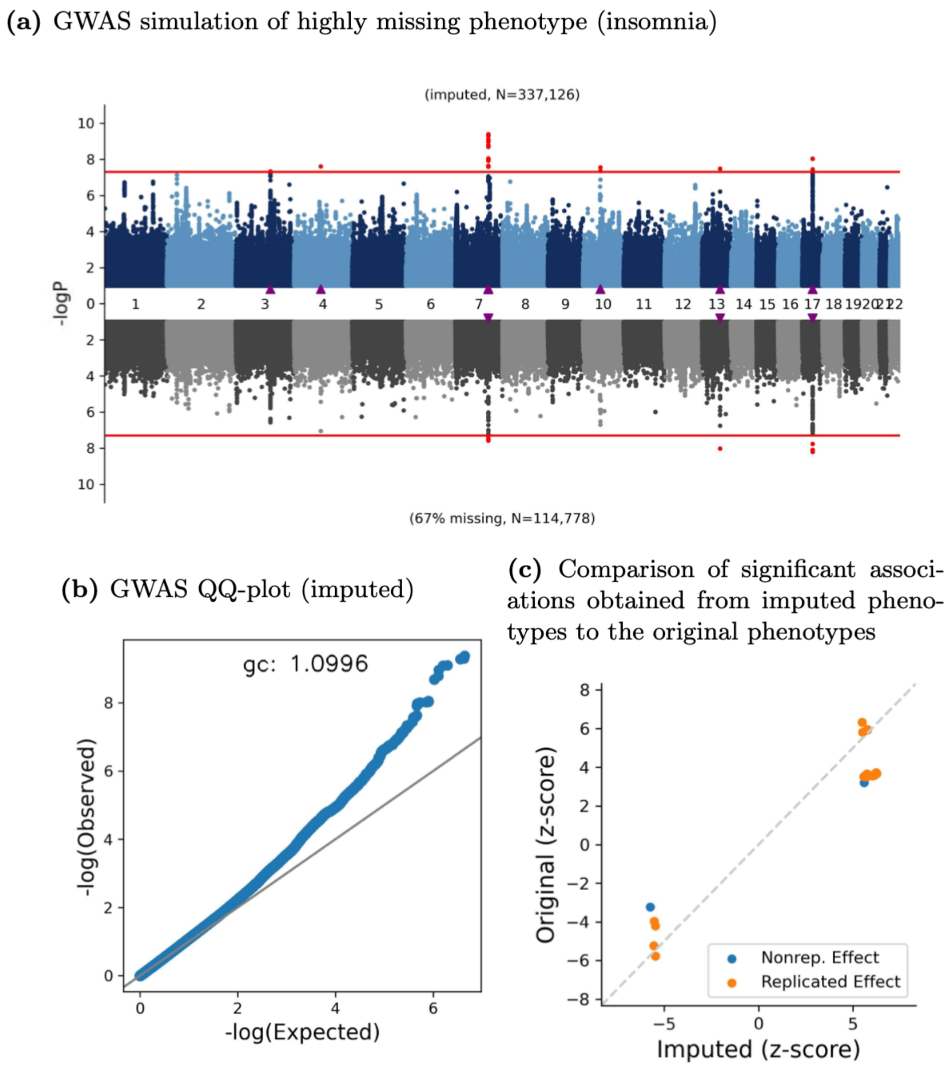


**Figure S3:** We simulated missing values (67% missingness) in a phenotype (insomnia at baseline measurement) that had low missingness and imputed missing values using AutoComplete. (a) GWAS for imputed alcohol consumption (top, N=337,126) compared to GWAS on the observed phenotype (bottom, N=112,297). Triangle markers indicate significantly associated loci ($p < 5 \times 10^{-8}$ indicated by the red horizontal line). The number of significant loci increased from 3 to 6 corresponding to an increase in the number of significantly associated SNPs from 10 to 28. (b) The corresponding QQ-plot of the GWAS augmented with imputed measurements is shown. (c) We attempted to test whether the significant associations obtained from the imputed phenotypes ("Imputed") were truly associated with the original phenotype (100% observed in data, "Original"). Z-scores of effect sizes for each experiment were plotted against one another. Significant associations in the imputed phenotype ($p < 5 \times 10^{-8}$) were determined as having a verifiable effect in the fully observed phenotype GWAS for $p < 5 \times 10^{-4}$ (accounting for the number of SNPs tested). Most (26 of 28) of the SNPs that were significantly associated in the imputed test could be verified as being strongly associated in the original test. The corresponding QQ-plot of the GWAS with imputed measurements for two tested phenotypes (alcohol consumption and insomnia) are shown in (c) and (d).

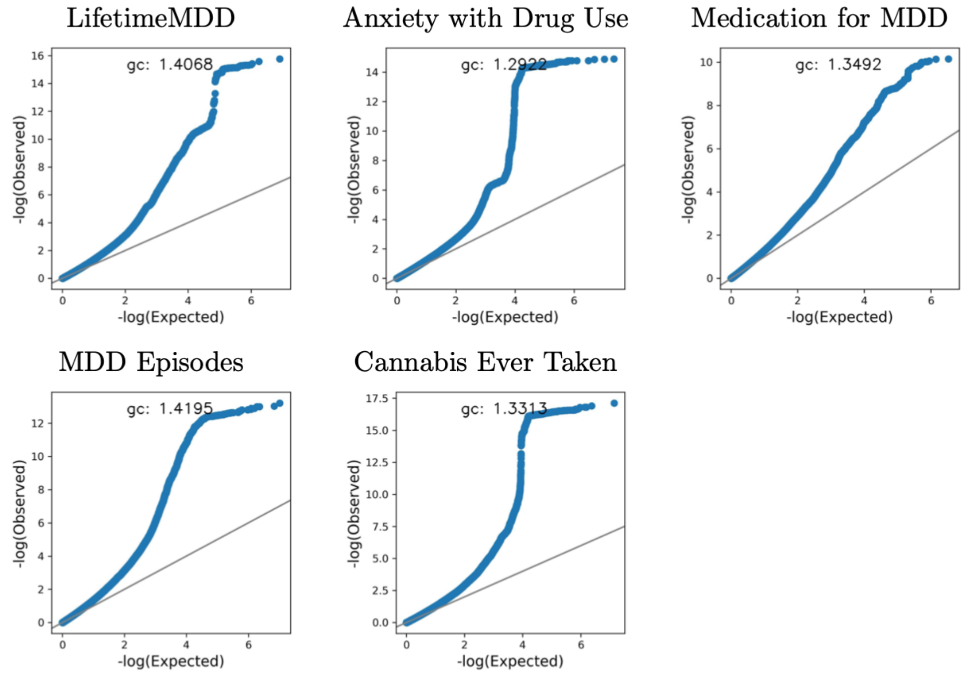


**Figure S4:** QQ-plots corresponding to GWAS of five highly missing phenotypes after imputation with AutoComplete.

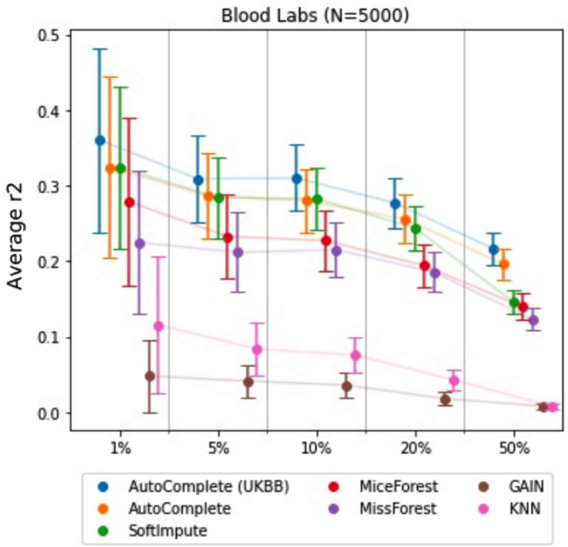


**Figure S5:** Imputation accuracy ( $r^2$ ) was evaluated for a smaller setting of N=5,000 randomly drawn subset of individuals from the blood labs dataset for which MissForest and MiceForest could be effectively fit. Accuracy obtained from fitting AutoComplete on the full training set of N=151,273 is also shown in blue (“AutoComplete (UKBB)”).

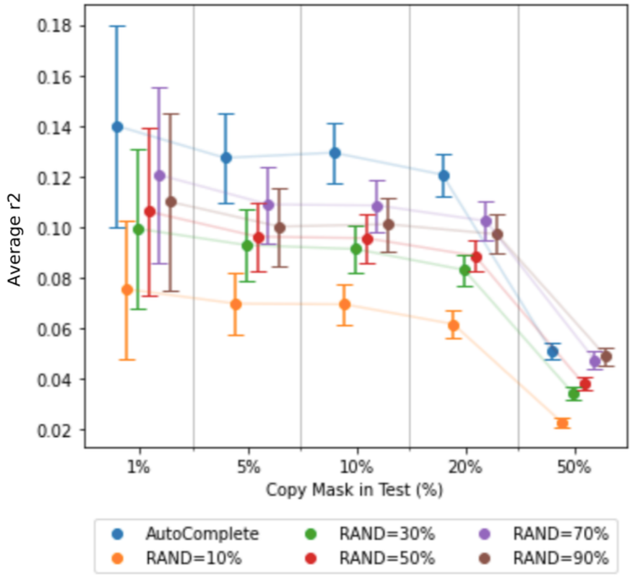


**Figure S6:** The contribution of copy-masking to our method was assessed in the context of training using only random masking of varying amounts with no copy-masking (RAND=10%~90%). Imputation accuracy ($r^2$) across phenotypes for the psychiatric disorder dataset was reported for each setting to determine overall performance. Imputation accuracy was measured for increasing percentages (1%~50%) of simulated missingness of the original data.

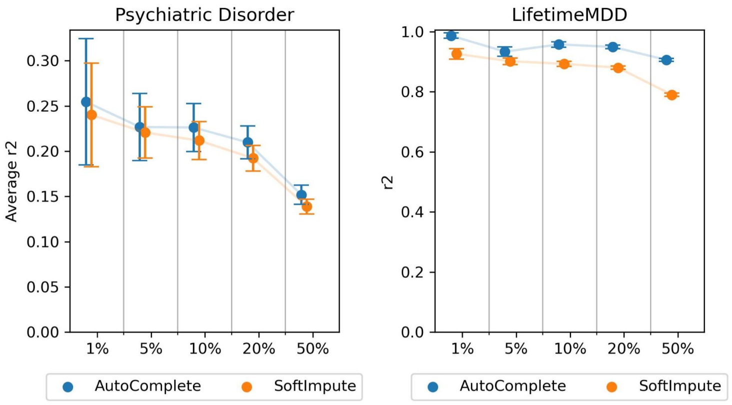


**Figure S7:** Evaluation of imputation accuracy using uniform masking instead of copy-masking for simulated missingness. Uniformly random masking was applied in increasing amounts as a percentage of observed data (1%~50%). AutoComplete and SoftImpute models tuned for the main experiments were reused and evaluated for the uniform missingness setting. We plot the accuracy averaged across phenotypes as well as the accuracy of one phenotype of interest (LifetimeMDD).
